# Identification of a Novel Retrieval-Dependent Memory Process in the Crab *Neohelice granulata*

**DOI:** 10.1101/2019.12.19.881128

**Authors:** Santiago A. Merlo, Maria J. Santos, Maria E. Pedreira, Emiliano Merlo

## Abstract

Fully consolidated associative memories may be altered by alternative retrieval dependent memory processes. While a brief exposure to the conditioned stimulus (CS) can trigger reconsolidation of the original memory, a prolonged CS exposure will trigger memory extinction. The conditioned response is maintained after reconsolidation, but is inhibited after extinction, presumably by the formation of a new inhibitory memory trace. In rats and humans, it has been shown that CS exposure of intermediate duration leave the memory in an insensitive or limbo state. Limbo is characterised by the absence of reconsolidation or extinction. Here we investigated the evolutionary conserved nature of limbo using a contextual Pavlovian conditioning (CPC) memory paradigm in the crab *Neohelice granulata*. In animals with fully consolidated CPC memory, systemic administration of the protein synthesis inhibitor cycloheximide after 1 CS presentation disrupted the memory, presumably by interfering with memory reconsolidation. The same intervention given after 320 CSs prevented CPC memory extinction. Cycloheximide had no behavioural effect when administered after 80 CS presentations, a protocol that failed to extinguish CPC memory. Also, we observed that a stronger CPC memory engaged reconsolidation after 80 CS instead of limbo, indicating that memory strength affects the parametrical conditions to engage either reconsolidation or limbo. Altogether, these results indicate that limbo is an evolutionary conserved memory process segregating reconsolidation from extinction in the number of CSs space. Limbo appears as an intrinsic component of retrieval dependent memory processing, with a key function in the transition from memory maintenance to inhibition.

**Author statement (CRediT Roles):** **Santiago A. Merlo:** Conceptualization, Data curation, Formal Analysis, Investigation, Software, Validation, Visualization, Writing – original draft, Writing – review & editing.

**Jimena Santos:** Investigation, Writing – review & editing.

**Maria Eugenia Pedreira:** Conceptualization, Funding acquisition, Project administration, Resources, Supervision, Validation, Writing – original draft, Writing – review & editing.

**Emiliano Merlo:** Conceptualization, Funding acquisition, Project administration, Resources, Supervision, Validation, Writing – original draft, Writing – review & editing.

## 1. Introduction

Paired presentations of an environmental stimulus that initially has no intrinsic biological significance (conditioned stimulus, CS), with a biologically relevant outcome (unconditioned stimulus, US), lead to the formation of an associative memory CS-US. Recently formed associative memories exist in a fragile state and are susceptible to behavioural or pharmacological disruption. However, through consolidation these memories become resistant to disruption (Dudai, Karni, and Born, 2015). For decades it was believed that fully consolidated memories remained unchangeable, but recent evidence indicate that memories are dynamic entities and their content or strength could be affected in a retrieval dependent manner (Reichelt and Lee, 2013).

In animals with a fully consolidated CS-US memory, presentation of the CS alone can trigger alternative memory processes, depending on the presentation parameters. On the one hand, a brief CS alone exposure will typically destabilise the memory, making it sensitive to amnestic agents, and trigger a restabilisation process that returns the memory to a stable state. This process called memory reconsolidation maintains the conditioned response towards the CS and constitutes an opportunity for updating the original CS-US memory, enabling changes in strength and content (Forcato, Rodriguez, and Pedreira, 2011; Forcato, Rodriguez, Pedreira, and Maldonado, 2010; Frenkel, Maldonado, and Delorenzi, 2005; Fukushima, Zhang, Archbold, Ishikawa, Nader, and Kida, 2014; Lee, 2010). On the other hand, a more prolonged CS exposure will more likely trigger memory extinction and a reduction of the conditioned response. Importantly, extinction does not erase the original memory, but relies on formation of a new associative trace between the CS and the absence of the US (CS-noUS) and concomitant decrease in CS-US memory expression (Bouton, 2004; Delamater and Westbrook, 2014; Pagani and Merlo, 2019). Hence, depending on the duration or number of CS alone events, in animals with fully consolidated CS-US memory retrieval could trigger alternative and behaviourally opposite processes (Lee, Milton, and Everitt, 2006; Pedreira and Maldonado, 2003). Increasing exposure to the CS alone shifts the neural system from maintenance to inhibition of the original memory engaging reconsolidation or extinction, respectively. Recently it was reported that these memory mechanisms are mutually exclusive, and that intermediate CS exposure sessions fail to engage either process. In auditory fear conditioned rats intermediate CS exposures engage a novel retrieval-dependent process which we have called ‘limbo’, characterised by insensitivity of the CS-US memory to amnestic treatments and a lack of extinction-specific molecular correlates (Merlo, Milton, and Everitt, 2018; Merlo, Milton, Goozee, Theobald, and Everitt, 2014). Limbo has also been documented for Pavlovian-conditioned contextual fear, and appetitive memories in rats (Cassini, Flavell, Amaral, and Lee, 2017; Flavell and Lee, 2013; Franzen, Giachero, and Bertoglio, 2019), as well as conditioned fear in humans (Kindt, Soeter, and Sevenster, 2014). There is little mechanistic information about limbo. Canonical plasticity-related mechanisms of reconsolidation and extinction, such as requirement of NMDA-type glutamate or GABA receptor activity, or activation of the kinase ERK1/2 in the basolateral amygdala, are not necessary (Franzen et al., 2019; Merlo et al., 2018; Merlo et al., 2014). Thus far, protein synthesis requirement during limbo has not been established. Considering the remarkable evolutionary persistence of most underlying cellular and molecular mechanisms of memory processes across species, in this work we will analyse limbo protein synthesis dependence using an associative memory paradigm in crabs.

Associative memory experiments in the crab *Neohelice granulata* have made important contributions towards better understanding the effects of retrieval on memory persistence. Besides documenting that reconsolidation and extinction are evolutionary conserved features, research in crabs indicates that triggering reconsolidation requires prediction error in the form of a mismatch between training and retrieval conditions or changes in the reinforcement ratio (López, Santos, Cortasa, Fernandez, Carbó Tano, and Pedreira, 2016; Pedreira, Perez-Cuesta, and Maldonado, 2004). The crab’s associative learning paradigm is based on the animal’s innate escape response, which is elicited by the presentation of an inescapable visual danger stimulus (VDS; an opaque rectangle moving horizontally above the animal). Repeated VDS presentations induce a change in behavioural response from escape to freezing (Pereyra, Gonzalez Portino, and Maldonado, 2000) and formation of an associative memory between the VDS (US) and a cue light (CS) presented above the animal (Fustiñana, Carbo Tano, Romano, and Pedreira, 2013). Escape response from trained animals at test, typically conducted at 24 hours or more, will be significantly lower than untrained controls, indicating the formation of an associative CS-US long-term memory. Animals with a fully consolidated so-called contextual Pavlovian conditioning (CPC) memory undergo reconsolidation when exposed to a brief CS alone trial, and extinction when exposed to a prolonged CS (Fustiñana et al., 2013). Both memory reconsolidation and extinction in the crab require protein synthesis, among other conserved neural mechanisms (Pedreira and Maldonado, 2003).

We tested the hypothesis that intermediate number of CS alone presentations will trigger limbo of the CPC memory. We predicted that in crabs with consolidated CPC memory, systemic administration of the protein synthesis inhibitor cycloheximide will: 1) disrupt the original CPC memory after exposure to a small number of CSs; 2) disrupt CPC extinction after a prolonged CS exposure session; and 3) have no behavioural effect after an intermediate CS exposure session. If so, our results will indicate that, not only is limbo an evolutionary conserved feature of retrieval-dependent memory processing but also, that it is protein synthesis independent. Additionally, we analysed whether a stronger CPC memory would require a higher number of CS presentations to enter limbo, in comparison with a less-strong ‘standard’ memory. In this regard, we predicted that animals trained with 30 CS-US trials would require a larger number of CS presentations to engage limbo compared to animals trained with 15 trials.

## 2. Materials & Methods

### 2.1 Animals

Adult male *Neohelice granulata* (formerly known as *Chasmagnathus granulatus*, Crustacea, Grapsidae) intertidal crabs, measuring 2.6-2.9 cm across the carapace and weighting ∼17 g, were collected from water <1 m deep in the estuarine coasts of San Clemente del Tuyú, Buenos Aires province, Argentina. Once transported to the laboratory, they were housed in plastic tanks (30 × 45 × 20 cm) filled to 0.5 cm depth with diluted (12‰, pH 7.4-7.6) marine water (Red Sea’s Coral Pro Salt) to a density of 20 crabs per tank. The holding room was maintained on a 12-h light-dark cycle (light on at 07:00). Temperature on both holding and experimental rooms was maintained at 22-24 °C. Experiments were carried out between the 3rd and the 10th day after the arrival of the animals. Each crab was used in only one experiment. Experimental procedures are in compliance with the National Institutes of Health Guide for Care and Use of Laboratory Animals (USA) and the Argentinean guidelines on the ethical use of animals in laboratory experiments. All efforts were made to minimize animal suffering and to reduce the number of animals used.

### 2.2 Experimental device

The experimental device has been described in detail elsewhere (Fustiñana et al., 2013). Briefly, the experimental unit was a bowl-shaped opaque container surrounded by a steep concave wall 12 cm high (23 cm top diameter and 9 cm floor diameter). The container was filled with marine water to a depth of 0.5 cm. The crab was placed in the container and could freely move inside but was not able to escape from it. The container could be illuminated from above or below by using an array of 3 LED bulbs (3mm, 1w each) and 10-W daylight lamp, respectively. A motor-operated screen (US, an opaque rectangular strip of 25.0 × 7.5 cm) was moved horizontally over the animal from left to right, and vice versa. The screen’s movements were cyclical. The screen displacements provoked the crab escape response. Each trial consisted of two successive cycles of screen movement. The experimental room had 20 experimental units separated by white walls, so that 20 crabs could be trained or tested independently and simultaneously. In each experiment, crabs and containers were assigned in a counterbalanced manner such that subjects belonging to each experimental group were included in every session. The experimental scheme was repeated until reaching the final number of animals for the experiment. Escape responses were video recorded at 10 Hz. Two days before the beginning of each experiment, animals were marked with a little round piece (0.5 cm diameter) of white ethylene-vinyl acetate glued to the centre of the carapace. Animal movement during training and testing trials was analysed by tracking x-y displacement of the white spots over time, using a customized image tracking software. A computer was used to programme trial sequences, illumination and duration, inter-trial intervals, and to monitor the experimental events.

### 2.3 Experimental procedure

All experimental groups started with 40 crabs. Between sessions, animals were housed individually in opaque plastic cylinders filled to depth of 0.5 cm with artificial sea water, inside dimly lit drawers and isolated from external stimulation. All behavioural sessions were preceded by 10 min of habituation to the experimental device with illumination from below. *Training session (Day 1)*. A ‘standard’ contextual Pavlovian conditioning (CPC) training procedure consisted of 15 trials, whereas a ‘strong’ training procedure consisted of 30 trials. Each trial lasted 27 s with above illumination (CS), and the US was presented during the last 9 s. During the intertrial interval (ITI; 153 s), the experimental unit was illuminated from below, which provoked a virtual change in the environmental features. All animals received one US presentation to evaluate individual reactivity. Untrained animals were kept in the experimental unit during the entire training procedure and were presented with the same pattern of light shift than trained animals, but without USs.

#### CS exposure session and drug administration (Day 2)

Twenty-four hours after training, crabs were placed in their respective training context. A specific amount of CS alone events (1, 40, 80, 160 or 320 presentations, depending on the experiment) were presented to each animal. Each CS presentation consisted of 27 s of above illumination (CS, ITI = 27 s). Immediately after, crabs were returned to their individual housing container. One hour later they were injected with either drug or vehicle solution.

#### Test session (Day 3)

Twenty-four hours after CS exposure, crabs returned to their training context and received a CS-US trial during which the escape response was measured.

### 2.4 Drugs and injection procedure

The protein synthesis inhibitor cycloheximide (CHX, Sigma-Aldrich C7698) (Pedreira, Dimant, Tomsic, Quesada-Allue, and Maldonado, 1995) was diluted in crustacean saline solution (450 mM NaCl, 15 mM CaCl_2_, 21 mM MgCl_2_, 10 mM KCl) (Hoeger and Florey, 1989) and administered systemically at a final dose of 2.35 µg/g. Fifty microlitres of vehicle (VHC) or drug solution were injected through the right side of the dorsal cephalothoraxic-abdominal membrane by means of a syringe needle fitted with a sleeve to control the depth of penetration to 4 mm, thus ensuring that the injected solution was released in the pericardial sac. Given that crabs lack an endothelial blood-brain barrier (Abbott, 1970), and that blood is distributed by a capillary system in the central nervous system (Sandeman, 1967), systemic injected drugs readily reach the brain.

### 2.5 Data analysis and drug effect evaluation

Memory retention was defined as a statistically significant lower conditioned response level on the testing session by the trained (TR) group, relative to its respective untrained (CT) group (i.e. both groups were injected with the same solution or exposed to the same number of CSs during Day 2). A lack of difference between a CT-TR pair was taken to indicate no CPC memory retention. Depending on experimental conditions, this could be due to amnesia or CPC memory extinction. A comparison between CT groups injected with either CHX or VHC was necessary to control for CHX side effects that may have affected response level at testing in a manner unrelated to the behavioural experience. In general, statistical analysis of test data included three *a priori* planned comparisons: CT-VHC vs. TR-VHC; CT-VHC vs. CT-CHX; and CT-CHX vs. TR-CHX. These comparisons were based on extensive prior experience indicating that spaced training with 15 or more trials results in memory retention, and on the prediction that the specific pharmacological manipulation should be free of non-mnemonic behavioural effects. If these two predictions were fulfilled the experiment will be valid to analyse the effect of the drug on memory. Escape responses are represented as mean ± SEM normalised to the respective vehicle control group (CT-VHC). Behavioural data were analysed using a one-way analysis of variance (ANOVA) with α (per comparison error rate) = 0.05, and *a priori* planned comparisons using IBM SPSS Statistics software (Version 22.0; Armonk, NY: IBM Corp). Effect sizes were calculated using partial eta square.

Given that some of our predictions about the engagement of limbo for intermediate number of CS presentations favoured the alternative hypothesis over the null hypothesis, we also analysed the data with Bayesian statistics. For each experiment we used Bayesian ANOVA followed by the same comparisons made in frequentist analysis using one-tailed Bayesian Independent Samples T-Tests (JASP, v0.10; JASP Team, 2019). BFs were interpreted using the categories proposed by Jeffreys (Jeffreys, 1961; Wetzels, Matzke, Lee, Rouder, Iverson, and Wagenmakers, 2011). BF_10_ represents the probability of the data to be explained by the alternative hypothesis (H_1_: at test, escape response in TR group is lower than CT) relative to the null hypothesis (H_0_: at test, escape response in TR group is similar to CT). For clarity, BF_10_ < 1 indicates that the null hypothesis was 1/BF_10_ more likely to explain the data, than the alternative hypothesis.

## 3. Results

### 3.1 A protein synthesis independent memory process segregates reconsolidation and extinction of CPC memory

Crabs with fully consolidated CPC memory will undergo reconsolidation or extinction when exposed to a brief (27 s) or prolonged (2h) CS alone event, respectively (Fustiñana et al., 2013). In order to test our main hypothesis, stating that an intermediate CS exposure session will trigger limbo in crabs with fully consolidated CPC memory, we conducted three independent experiments. Considering that both memory reconsolidation and extinction are proteins synthesis dependent mechanisms, we used post-retrieval systemic injections of cycloheximide as a tool to reveal the dominant memory process after different CS exposure sessions.

We first evaluated the effect of cycloheximide after a single CS alone presentation. Animals were either trained with 15 CS-US trials (TR group) or remained in the training context for a similar duration (CT group), during Day 1. Twenty-four hours later, all animals were exposure to one CS alone event. One hour later, both CT and TR groups were divided and injected with either vehicle (CT-VHC, TR-VHC) or cycloheximide solution (CT-CHX, TR-CHX) before returning to their housing container. On Day 3, CPC memory was evaluated by presenting all animals with a single CS-US trial (Figure 1A, n_CS_ = 1). One-way ANOVA of the escape response during test showed a main effect of group (F _(3, 105)_ = 2.90, *p* = 0.04, η^2^ = 0.08). As expected, the vehicle injected CT-TR pair showed memory retention, with the TR group expressing a lower escape response compared to CT (CT-VHC vs TR-VHC: *t* _(105)_ = 2.75, *p* = 0.007, η^2^ = 0.12). Escape response in CT groups were similar (CT-VHC vs CT-CHX: *t* _(105)_ = 0.37, *p* = 0.71, η^2^ = 0.003), indicating there were no non-specific effects of CHX at test. Cycloheximide injected CT-TR groups showed similar escape response at test (CT-CHX vs TR-CHX: *t* _(105)_ = 0.48, *p* = 0.63, η^2^ = 0.005), indicating the lack of CPC memory retention in TR-CHX animals (Figure 1B). Bayes analysis on the vehicle injected CT-TR pair indicated that the alternative hypothesis (i.e. escape response in TR animals is different to CT animals) is 4.08 times more likely than the null hypothesis (CT-VHC vs TR-VHC: BF_10_ = 4.08). For the cycloheximide injected CT-TR pair the null hypothesis (i.e. escape response in TR animals was similar to CT animals) is 3.1 times more likely than the alternative hypothesis (CT-CHX vs TR-CHX: BF_10_ = 0.32). A similar outcome was found for comparison of both CT groups (CT-VHC vs CT-CHX: BF_10_ = 0.29). Hence, both frequentist and Bayesian analysis support the conclusion that crabs with fully consolidated CPC memory presented with 1 CS alone engage memory labilisation-reconsolidation.

**Figure 1:**
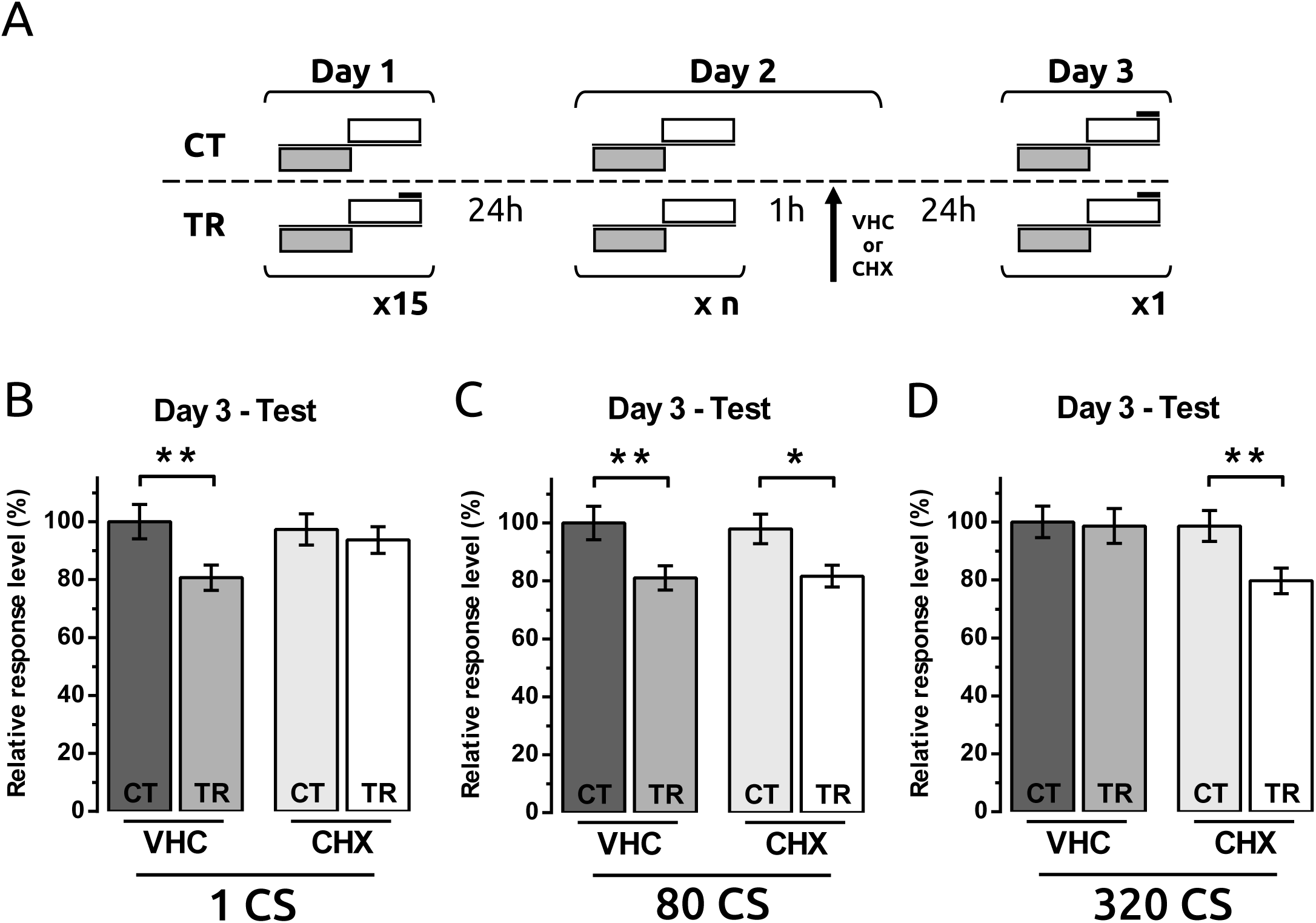
Effect of systemic cycloheximide administration on alternative retrieval-dependent memory processing in the crab *Neohelice granulata*. **A.** Experimental design: On Day 1 crabs were trained with 15 trials (ITI = 153 sec, TR groups), or remained in the context without US presentation (CT groups). On Day 2 crabs were presented with either 1, 80 or 320 CS alone trials (**B, C** or **D**, respectively). One hour later animals received a systemic injection of cycloheximide or vehicle solution, forming four experimental groups per CS condition. On Day 3 CPC memory retention was evaluated by presentation of a single CS-US trial. **B-D.** Mean relative escape response level (± SEM) at test (Day 3) are shown for each CS exposure at Day 2 condition. Group sizes: **B**, CT-VHC = 30, TR-VHC = 29, CT-CHX = 25, TR-CHX = 25; **C**, CT-VHC = 38, TR-VHC = 38, CT-CHX = 33, TR-CHX = 34; **D**, CT-VHC = 38, TR-VHC = 40, CT-CHX = 39, TR-CHX = 39. \**p* < 0.05. \*\**p* < 0.01.

Next we analysed the effect of post-retrieval cycloheximide after exposure to an intermediate number of CS presentations. The experimental design was similar to the previous experiment, but on Day 2 animals were exposed to 80 CS presentations, in the absence of USs (Figure 1A, n_CS_ = 80). Analysis of the escape response during test showed a main effect of group (ANOVA: F _(3, 139)_ = 4.61, *p* = 0.004, η^2^ = 0.09). Both vehicle and cycloheximide injected CT-TR groups showed memory retention (CT-VHC vs TR-VHC: *t* _(139)_ = 2.89, *p* = 0.005, η^2^ = 0.10; CT-CHX vs TR-CHX: *t* _(139)_ = 2.33, *p* = 0.021, η^2^ = 0.08), with no non-specific effect of CHX (CT-VHC vs. CT-CHX: *t* _(139)_ = 0.30, *p* = 0.77, η^2^ = 0.001; Figure 1C). Bayesian analysis indicated that in either vehicle or cycloheximide injected CT-TR pairs the alternative hypothesis was 4.8 and 4.1 more likely than the null hypothesis, respectively (CT-VEH vs TR-VEH: BF_10_ = 4.82; CT-CHX vs TR-CHX: BF_10_ = 4.08). The null hypothesis was 4 times more likely when comparing both CT groups (CT-VHC vs CT-CHX: BF_10_ = 0.25). These results indicate that 80 CSs were insufficient to produce extinction, but equally failed to render the CPC memory labile and sensitive to cycloheximide amnestic effect. Thus, CPC memory in crabs exposed to an intermediate number of CSs enters limbo.

In order to confirm that CPC memory extinguishes with larger CS exposure we run an additional experiment. Crabs were trained as before, but on Day 2 they were exposed to 320 CS alone presentations before vehicle or cycloheximide systemic administration (Figure 1A, n_CS_ = 320). An ANOVA on escape responses during test showed a main effect of group (F _(3, 152)_ = 3.32, *p* = 0.022, η^2^ = 0.06). Planned comparisons showed that there was no memory retention in the vehicle injected TR group (CT-VHC vs. TR-VHC: *t* _(152)_ = 0.18, *p* = 0.86, η^2^ = 0.0004) indicating the extinction of the conditioned response by 320 CSs alone presentations. Cycloheximide injected TR group showed a lower escape response at test compared with the CT group (CT-CHX vs. TR-CHX: *t* _(152)_ = 2.51, *p* = 0.013, η^2^ = 0.08), indicating extinction disruption due to protein synthesis inhibition after 320 CS presentations. Control groups showed a similar escape response (CT-VHC vs. CT-CHX: *t* _(152)_ = 0.18, *p* = 0.86, η^2^ = 0.0004), discarding non-specific effects of the drug at test (Figure 1D). Bayesian analysis showed that in the vehicle injected pair the null hypothesis is 4.2 more likely than the alternative hypothesis (CT-VHC vs TR-VHC: BF_10_ = 0.24), whereas in the cycloheximide injected pair the alternative hypothesis was 5.6 times more likely (CT-CHX vs TR-CHX: BF_10_ = 5.56). The null hypothesis was 4.2 times more likely when comparing both CT groups (CT-VHC vs CT-CHX: BF_10_ = 0.24).

Next we evaluated whether retrieval was necessary, but not sufficient, for cycloheximide amnestic effect over CPC memory. The experiment included three CT-TR pair of groups. On Day 2 two of the CT-TR pairs were exposed to either 1 or 80 CS presentations, whereas the third pair remained in their home container (non-reactivated pair, NR). One hour after CS exposure all animals received systemic cycloheximide injections (Figure 2A). At test, there was a main effect of group on escape responses (ANOVA: F _(5, 207)_ = 3.28, *p* = 0.007, η^2^ = 0.07). Planned comparisons showed that cycloheximide had no effect on CPC memory retention in the non-reactivated training group (CT-NR vs TR-NR: *t* _(207)_ = 2.61, *p* = 0.01, η^2^ = 0.09). There was no evidence of memory retention in the training group exposed to 1 CS and injected with the drug one hour later (CT-1CS vs TR-1CS: *t* _(207)_ = 0.73, *p* = 0.47, η^2^ = 0.007). As in our previous experiment, trained animals exposed to 80 CSs and injected with the drug showed memory retention (CT-80CS vs TR-80CS: *t* _(207)_ = 2.37, *p* = 0.02, η^2^ = 0.08). Bayesian analysis showed that in the non-reactivated or the 80 CS groups, the alternative hypothesis was 3 or 3.9 times more likely than the null hypothesis, respectively (CT-NR vs TR-NR: BF_10_ = 3.01; CT-80CS vs TR-80CS: BF_10_ = 3.89). For crabs exposed to 1 CS and injected with cycloheximide, the escape data at test was 3.2 times more likely to be explained by the null hypothesis (CT-1CS vs TR-1CS: BF_10_ = 0.31; Figure 2B).

**Figure 2:**
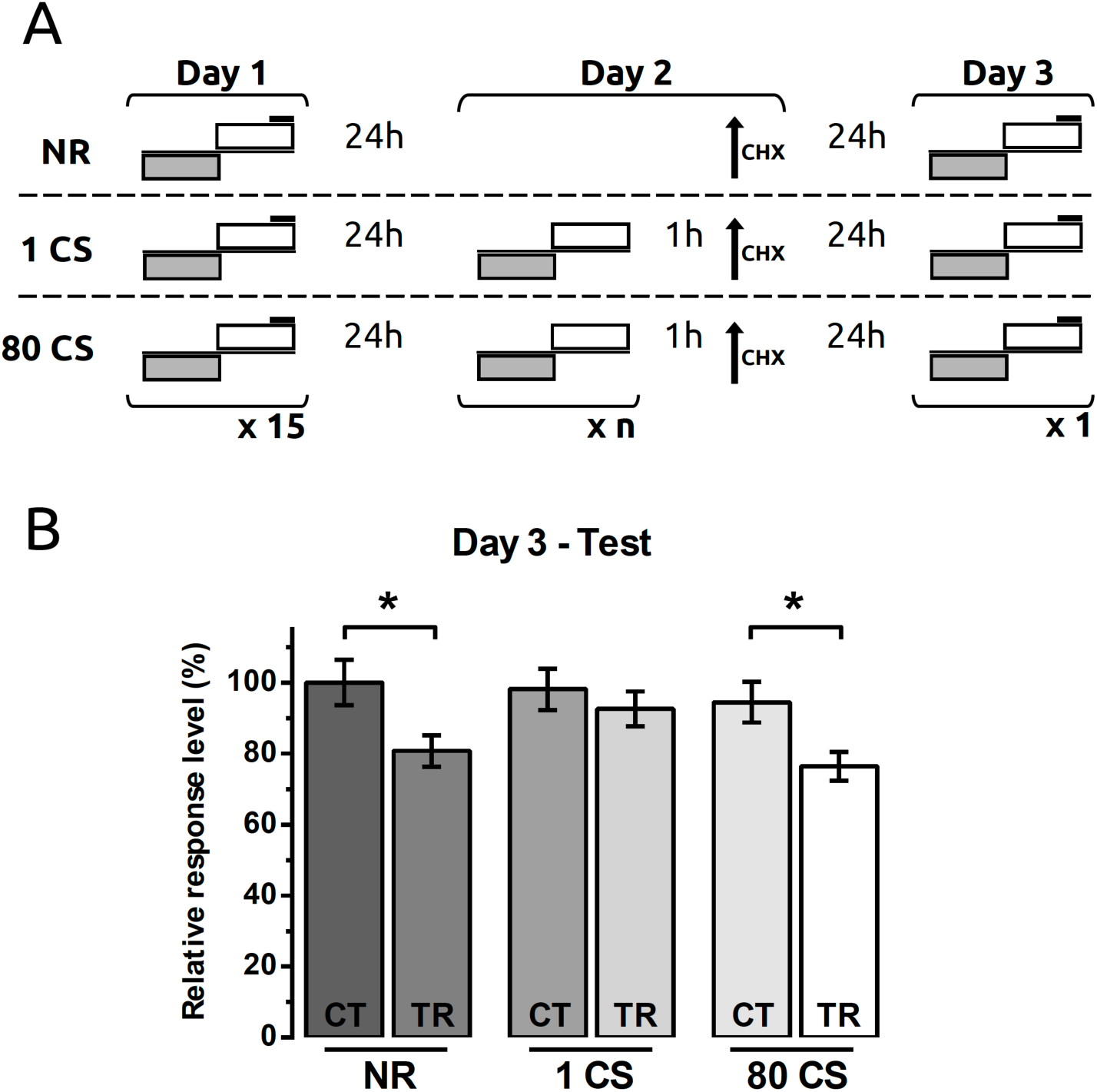
Effect of cycloheximide on CPC memory processing induced by 1 or 80 CS presentations. **A.** Experimental design: On Day 1 crabs were trained with 15 trials (ITI = 153 sec, TR groups), or remained in the context without US presentation (CT groups). On Day 2 crabs were presented with either 1 or 80 CS alone trials or remained in their home containers (NR groups). One hour later all groups received a systemic injection of cycloheximide. On Day 3 CPC memory retention was evaluated by presentation of a single CS-US trial. **B.** Mean relative escape response level (± SEM) at test (Day 3) are shown for each CS exposure at Day 2 condition. Group sizes: CT-NR = 38, TR-NR = 36, CT-1CS = 34, TR-1CS = 36, CT-80CS = 34, TR-80CS = 35. \**p* < 0.05.

Once more, we observed that cycloheximide disrupted CPC memory after 1 CS presentation in a retrieval dependent manner but has no effect on CPC memory when administered after 80 CS presentations. Even though CPC memory retrieval is a necessary condition for cycloheximide to exert an amnestic effect, it is not sufficient. Retrieval duration critically affects memory sensitivity to protein synthesis inhibition.

In order to gain a better insight on memory lability along increasing duration of CS exposure, we evaluated the effect of cycloheximide on memory performance after 40 or 160 CS alone presentation sessions. Cycloheximide disrupted CPC memory when administered after 40 CSs, indicating that this number of CSs also triggers memory labilisation-reconsolidation. Conversely, there was no effect of cycloheximide administered after 160 CS presentation (Supplementary Figure 1). After either 80 or 160 CS presentations, the original CPC memory was insensitive to CHX, and also non extinguished, strongly suggesting limbo engagement by these reminders.

These results not only confirmed the protein synthesis dependency of reconsolidation and extinction of CPC memory in crabs, but also shows for the first time that invertebrate memory processing also engages limbo after an intermediate number of CS presentations.

### 3.2 The boundary between reconsolidation and limbo depends on memory strength

In the previous experiment we established that a standard CPC memory enters limbo after an 80 CS long reminder session. We next examined whether the number of CSs necessary to engage limbo were dependent on memory strength.

We first evaluated the engagement of reconsolidation by presentation of a single CS alone event (Figure 3A, n_CS_ = 1). Crabs were trained with a ‘strong’ training session consisting of 30 CS-US trials (TR groups) or remained in the training environment for a similar amount of time (CT groups). Twenty-four hours later, animals were exposed to one CS trial and received an injection of vehicle or cycloheximide solution one hour later. An ANOVA on the escape responses during the test session at Day 3 showed a main effect of group (F _(3, 143)_ = 6.30, *p* < 0.001, η^2^ = 0.12). Planned comparisons showed memory retention for the vehicle injected TR group (CT-VHC vs. TR-VHC: *t* _(143)_ = 4.12, *p* < 0.001, η^2^ = 0.19), but not for the cycloheximide treated TR group (CT-CHX vs. TR-CHX: *t* _(143)_ = 1.37, *p* = 0.17, η^2^ = 0.03). Escape responding was similar for CT groups (CT-VHC vs. CT-CHX: *t* _(143)_ = 1.40, *p* = 0.16, η^2^ = 0.03), indicating no non-specific effect of the drug treatment at test (Figure 3B). Bayesian analysis indicated that in the vehicle injected CT-TR pair, the alternative hypothesis was 434 times more likely than the null hypothesis (CT-VHC vs TR-VHC, BF_10_ = 434.07), whereas for the cycloheximide injected pair, the data were 2 times more likely explained by the null hypothesis (CT-CHX vs TR-CHX, BF_10_ = 0.5). The null hypothesis was 1.9 times more likely when comparing both CT groups (CT-VHC vs CT-CHX, BF_10_ = 0.53).

**Figure 3:**
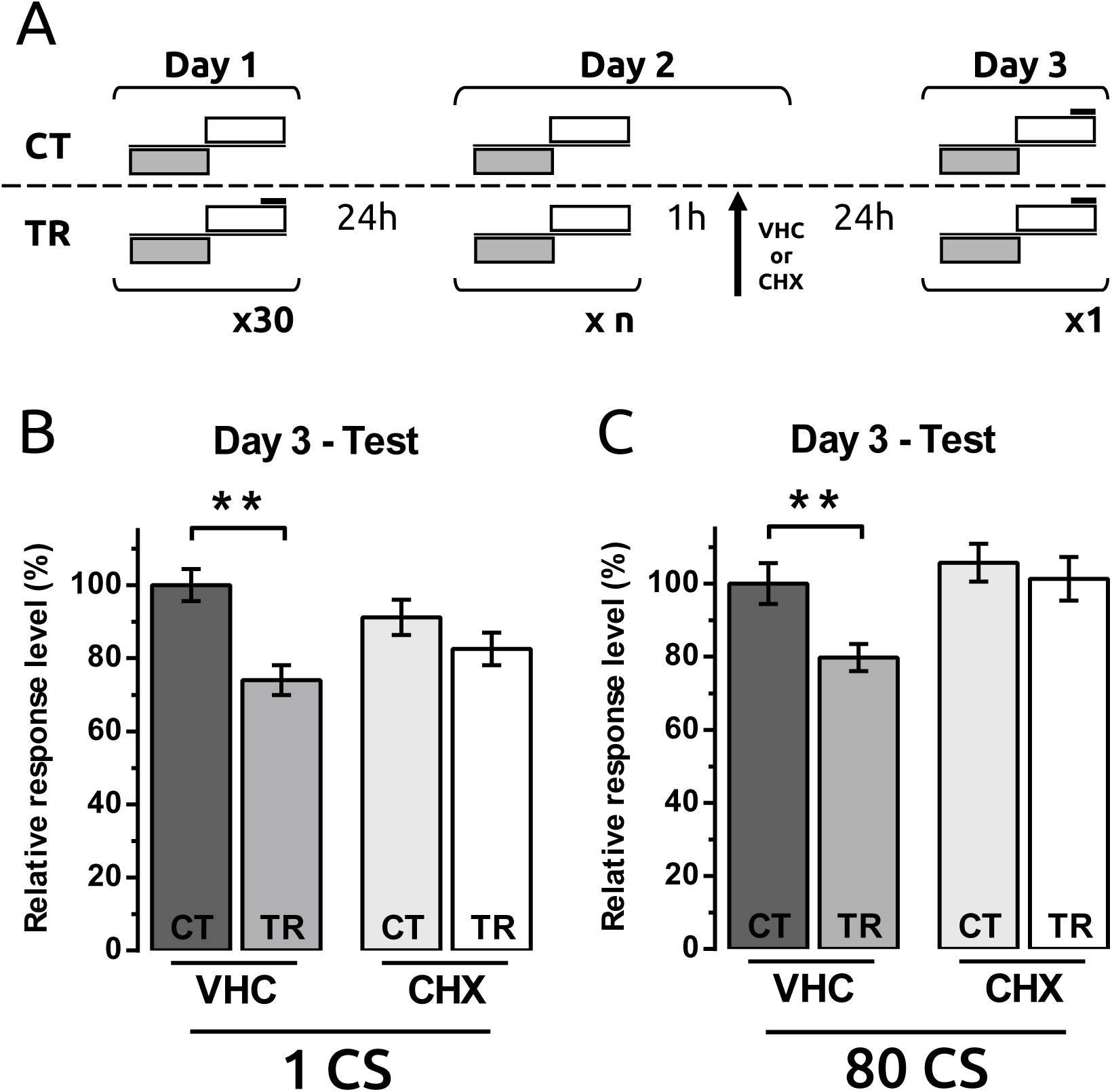
Effect of CPC memory strength on limbo engagement. **A.** Experimental design: same as in Figure 1, but the training session on Day 1 had 30 CS-US trials. On Day 2 animals were presented with either 1 or 80 CS alone trials. **B-C.** Mean relative escape response level (± SEM) at test (Day 3) are shown for each CS exposure at Day 2 condition. Group sizes: **B**, CT-VHC = 36, TR-VHC = 37, CT-CHX = 37, TR-CHX = 37; **C**, CT-VHC = 38, TR-VHC = 35, CT-CHX = 37, TR-CHX = 39. \*\**p* < 0.01.

Next we evaluated the effect of cycloheximide after exposure to 80 CSs, a manipulation that did not affect the conditioned response if crabs were trained with the standard protocol (Figure 1C). Animals were trained with 30 CS-US trials as before. Twenty-four hours later, all animals were presented with 80 CS alone trials and injected with either vehicle or cycloheximide solutions one hour later (Figure 3A, n_CS_ = 80). An ANOVA on the escape response level during test (Day 3) revealed a main effect of group (F _(3, 144)_ = 4.70, *p* = 0.004, η^2^ = 0.09). We observed memory retention for the TR-VEH group (CT-VHC vs. TR-VHC: *t* _(145)_ = 2.72, *p* = 0.007, η^2^ = 0.09), but amnesia for the TR-CHX group (CT-CHX vs. TR-CHX: *t* _(145)_ = 0.60, *p* = 0.55, η^2^ = 0.005). Both CT groups showed similar escape responses at test (CT-VHC vs. CT-CHX: *t* _(145)_ = 0.77, *p* = 0.44, η^2^ = 0.008), indicating no non-specific effect of the drug (Figure 3C). Bayesian analysis indicates that the alternative hypothesis is 9.4 times more likely in the vehicle injected CT-TR pair (CT-VHC vs TR-VHC, BF_10_ = 9.36), whereas the null hypothesis is 3.7 times more likely to explain the data for the cycloheximide injected pair (CT-CHX vs TR-CHX, BF_10_ = 0.27). The null hypothesis was 3.3 times more likely when comparing both CT groups (CT-VHC vs CT-CHX, BF_10_ = 0.3). These results show that a stronger CPC memory engages reconsolidation after either a single or 80 CS presentations, suggesting that the experimental parameters to trigger limbo are affected by memory strength such that stronger memories require a larger CS exposure session to engage limbo, with no effect on reconsolidation engagement by 1 CS presentation.

## 4. Discussion

In the present study we analysed the evolutionary conserved nature of limbo, a recently discovered retrieval dependent memory phase that segregates the processes of memory reconsolidation and extinction in the CS exposure domain. In particular, we evaluated whether in crabs with fully consolidated memory an intermediate re-exposure session would engage limbo, characterised by the absence of reconsolidation or extinction processes. Using post-retrieval systemic administration of cycloheximide as an amnestic agent, we observed that crabs with fully consolidated contextual Pavlovian conditioning (CPC) memory engaged memory reconsolidation after 1 or 40 CS alone events. To the contrary, presentation of 320 CSs triggered CPC memory extinction. Exposure to intermediate number of CSs (80 or 160) not only rendered the original CPC memory immune to cycloheximide, but also failed to extinguish it. This pattern of results clearly indicates that in crabs, reconsolidation and extinction are mutually exclusive processes, triggered by extreme reminder conditions. These alternative and opposing memory mechanisms are segregated in the CS exposure domain by an insensitive, limbo phase. Also, we observed that a stronger CPC memory requires more than 80 CSs to engage limbo, suggesting that memory strength is a boundary condition for limbo. These data indicate that limbo is present in invertebrates and independent of *de novo* protein synthesis, and that the absolute number of CSs that engage limbo is affected by memory attributes. We propose a transition model for retrieval dependent memory processing in crabs, with the processes of reconsolidation, limbo and extinction being engaged exclusively and sequentially as the number of CS presentations to animals with fully consolidated memory increases (Figure 4).

**Figure 4:**
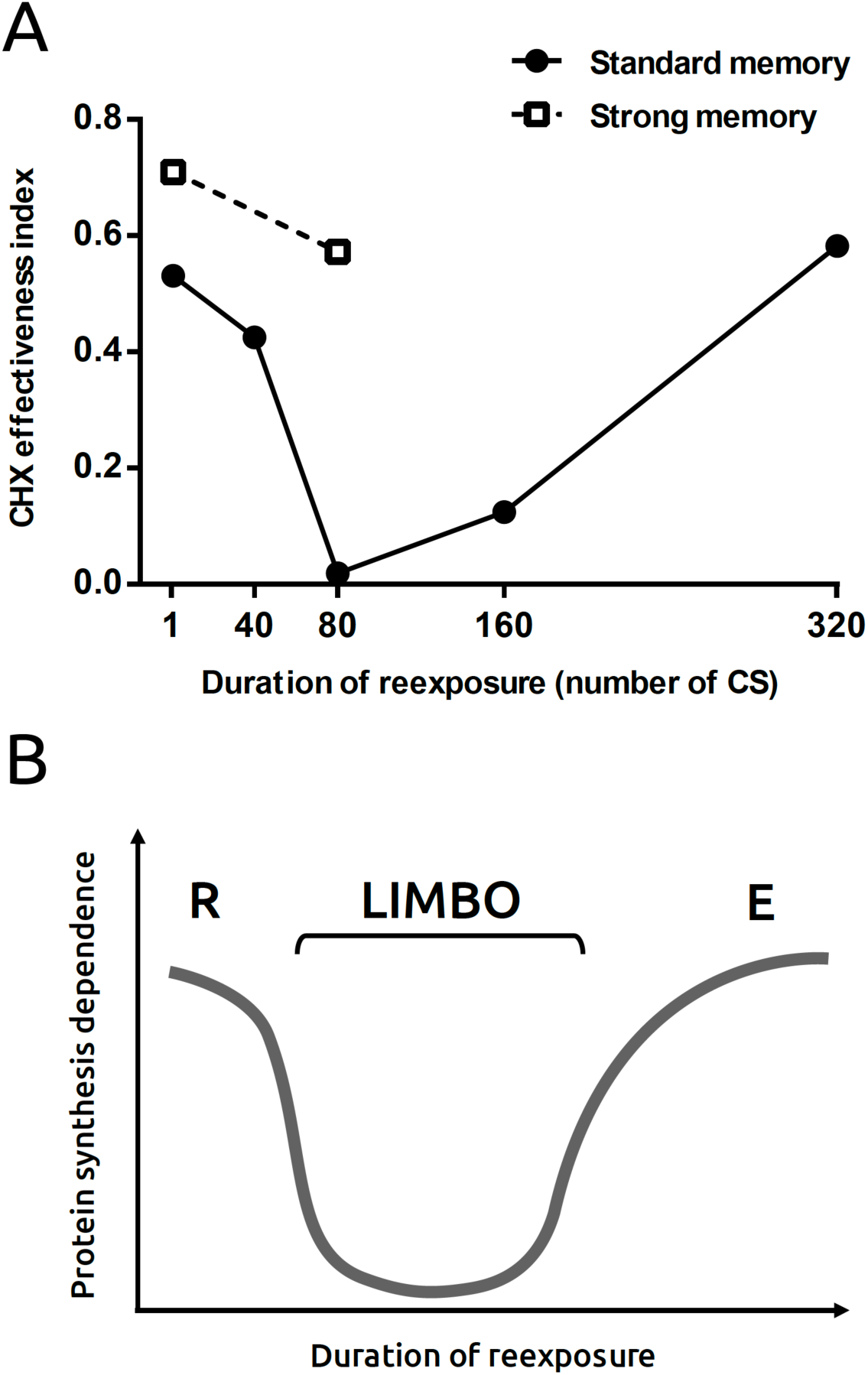
Alternative retrieval dependent memory processes in *Neohelice granulata*. **A.** Graphical representation of cycloheximide effect on memory after various reminder sessions. CHX effectiveness index is calculated as the absolute difference between effect size (Cohen’s d) in VHC groups and effect size in CHX groups during test. The graph shows CHX effectiveness index for different number of CS alone events from the experiments presented above. **B.** Schematic representation of protein synthesis dependency on alternative retrieval dependent memory process along the CS space. R: memory reconsolidation. L: memory in limbo. E: memory extinction.

### Limbo as an intrinsic component of retrieval dependent memory processing across animals

Limbo has been previously described as a retrieval dependent memory phase in vertebrates, affecting associative memories established with aversive and appetitive reinforcement. Our findings of alternative retrieval-dependent memory processes are not unprecedented and similar effects can be found across experimental paradigms and species. Blockade of β-adrenoreceptors by propranolol disrupts an associative fear memory in humans when administered after 2 CS presentations, whereas it has no behavioural effect after 4 CS presentations (Sevenster, Beckers, and Kindt, 2014). In cued fear conditioned rats, NMDA type glutamate receptor (NMDAR) antagonist MK-801 disrupts either reconsolidation or extinction when administered in conjunction with 1 or 10 CS presentations, respectively, but has no effect when given before 4 CSs. Similarly, NMDAR partial agonist D-cycloserine positively modulated the dominant memory process induced by 1 or 10 CS presentations but had no behavioural effect when given before 4 CSs (Merlo et al., 2014). Contextual fear conditioning in rats also shows a retrieval dependent insensitive phase with intermediate reminder durations. In this case, limbo was found using GABA enhancer midazolam (Alfei, Ferrer Monti, Molina, Bueno, and Urcelay, 2015; Franzen et al., 2019) or NMDAR antagonist MK-801 (Cassini et al., 2017). Moreover, limbo was also reported in rats lever pressing for food pellets. Systemic administration of MK-801 disrupted food seeking behaviour or its extinction in conjunction with a short (10 lever presses without food) or prolonged (50 lever presses) reminder session, respectively. However, MK-801 administration had no effect when given before an intermediate duration session (30 lever presses) (Flavell and Lee, 2013). In medaka fish, administration of an amnestic agent that disrupts reconsolidation or extinction under extreme reminder conditions had no behavioural effect when administered after an intermediate number of CSs (Eisenberg, Kobilo, Berman, and Dudai, 2003).

Finding limbo in crabs using cycloheximide injections, not only shows for the first time that it is an evolutionary conserved feature, but also that it is immune to the ‘gold standard’ amnestic manipulation in experimental neurobiology, inhibition of protein synthesis. This evidence strongly suggests limbo constitutes an intrinsic property of memory processing present along diverse taxa within the Animal Kingdom, from invertebrates to mammals. This common trait across phyla could be provided by common homologous circuits (Strausfeld and Hirth, 2013) supporting adaptive selection of alternative behaviours. Moreover, its protein synthesis independence sets limbo clearly apart in terms of neural mechanisms supporting both memory reconsolidation, extinction, and even consolidation. Limbo contribution to memory persistence or inhibition remains to be determined.

### Limbo boundary conditions

A close analysis of the reminder structure that leads to limbo in different memory paradigms highlights some key properties. Similar to memory reconsolidation and extinction processes, limbo engagement requires a reminder with prediction error, but of intermediate duration or number of events. Unlike memory extinction, limbo does not affect conditioned performance. Also, in crabs we observed that the sheer number of CS presentations required to trigger limbo is considerably larger to the number reported in other memory paradigms, suggesting that this variable is not important in isolation. Notably, in humans, limbo was induced after only 4 CS presentations (Sevenster et al., 2014), whereas in crabs it was observed after either 80 or 160 CS presentations. The ratios between CSs to engage limbo to CS-US trials at training are also variable across species, with the crab experiments requiring a ratio between 6 and more than 10, while other paradigms showed ratios of as little as 2 (Merlo et al., 2014; Sevenster et al., 2014). Moreover, limbo is not exclusively engaged in associative memories using discrete CSs as in our experiments; contextual fear memories, where environmental context serves as the CS, show that intermediate exposure duration also engage limbo (Cassini et al., 2017).

Here we discovered that limbo engagement depends on memory strength. Previous reports showed that memory strength affects the CS exposure required to trigger either memory reconsolidation or extinction. Stronger associative memories require longer CS alone exposure to enter reconsolidation or extinction, compared to a standard memory (Baumgartel, Genoux, Welzl, Tweedie-Cullen, Koshibu, Livingstone-Zatchej, Mamie, and Mansuy, 2008; Suzuki, Josselyn, Frankland, Masushige, Silva, and Kida, 2004). In this study, we observed that memory strength alters the parametrical conditions to engage reconsolidation and limbo. A stronger memory, produced by doubling the training duration from 15 to 30 CS-US trials, changed the efficacy of 80 CS presentations to engage limbo to memory reconsolidation (Figure 3). In contrast to previous observations showing a right-shift on reconsolidation engagement for stronger memories, reconsolidation of the stronger CPC memory was still engaged by 1 CS presentation. Thus, CPC memory strength in crabs modulates the upper limit of the CS exposure domain capable of inducing memory reconsolidation, without apparent effects on the lower limit. These observations reinforce the idea that sheer number of CS alone presentations is not a crucial parameter determining memory reconsolidation or limbo engagement. Altogether, these observations strongly suggest that memory strength is a boundary condition for the three alternative retrieval dependent memory processes: reconsolidation, limbo and memory extinction.

Finding that limbo is an essential part of retrieval dependent memory processes in invertebrates has both theoretical and practical implications. Even though the psychobiological function of limbo remains unclear, its evolutionary constancy suggests a putative role on memory persistence control. Limbo engagement might be necessary as a functional segregation between alternative and opposing memory processes, terminating reconsolidation and preparing the system for extinction. Clearly, more research will be necessary to answer these questions. Our work opens the possibility to explore limbo using a variety of experimental advantages and tools (e.g. simpler brains, easy access to neurons and networks, genetic manipulations) that are only available in invertebrates and can probe essential to reveal the intricate relationship of brain processes underlying persistence or inhibition of retrieved memories.

---

**Supplementary Figure 1:**
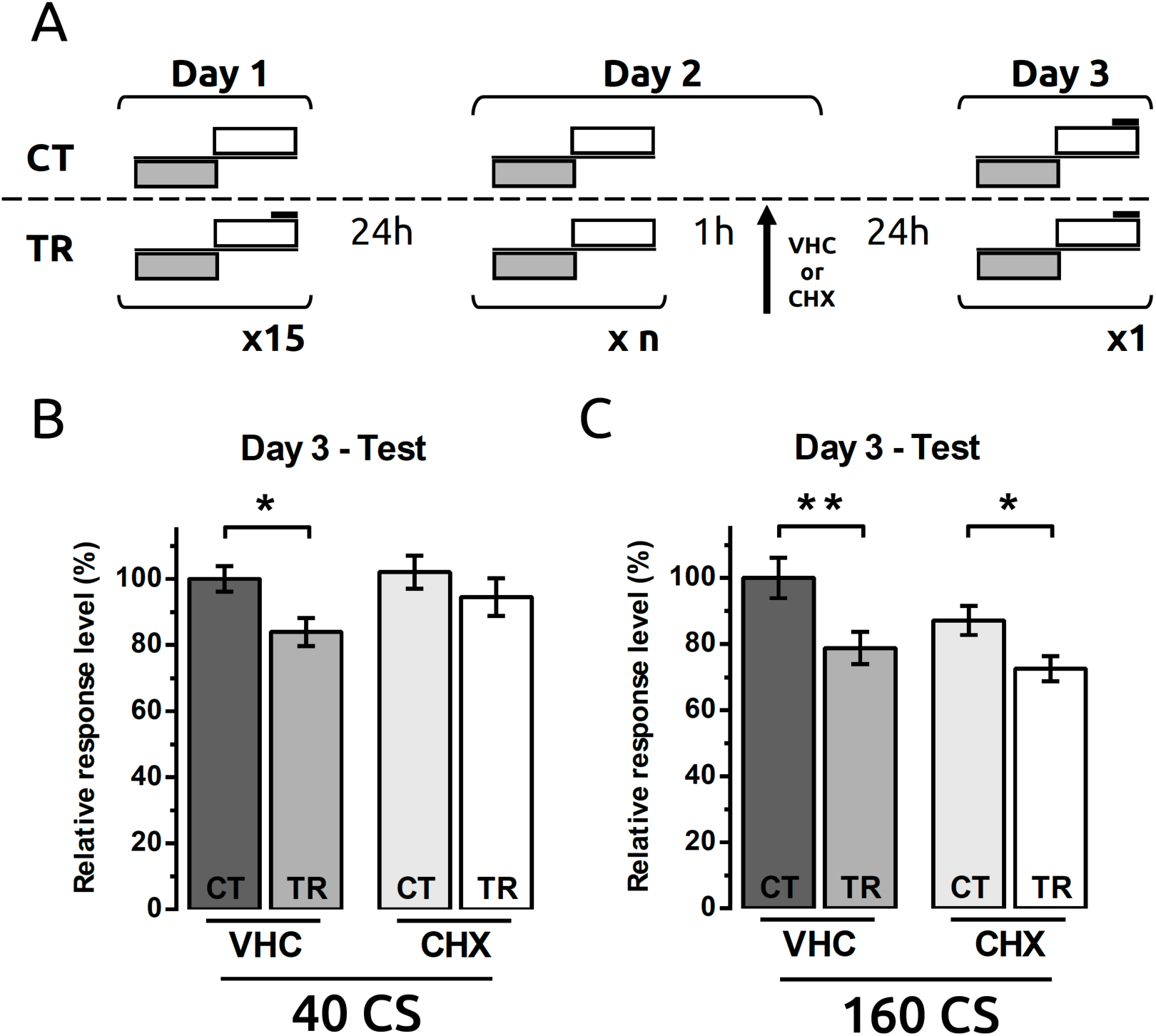
Effect of cycloheximide on CPC memory processing induced by 40 or 160 CS presentations. **A.** Experimental design: On Day 1 crabs were trained with 15 trials (ITI = 153 sec, TR groups), or remained in the context without US presentation (CT groups). On Day 2 crabs were presented with either 40 or 160 CS alone trials (**B** or **C**, respectively). One hour later animals received a systemic injection of cycloheximide or vehicle solution, forming four experimental groups per CS condition. On Day 3 CPC memory retention was evaluated by presentation of a single CS-US trial. **B-C.** Mean relative escape response level (± SEM) at test (Day 3) are shown for each CS exposure at Day 2 condition. **B.** Group sizes: CT-VHC = 36, TR-VHC = 36, CT-CHX = 35, TR-CHX = 37. ANOVA of escape response during test session: F _(3, 140)_ = 2.89, *p* = 0.04, η^2^ = 0.06). Planned contrasts: CT-VHC vs TR-VHC, *t* _(140)_ = 2.39, *p* = 0.02, η^2^ = 0.08; CT-VHC vs CT-CHX, *t* _(140)_ = 0.31, *p* = 0.76, η^2^ = 0.001; CT-CHX vs TR-CHX, *t* _(140)_ = 1.13, *p* = 0.26, η^2^ = 0.02. Bayesian analysis: CT-VHC vs TR-VHC, BF_10_ = 6.41; CT-VHC vs CT-CHX, BF_10_ = 0.26; CT-CHX vs TR-CHX, BF_10_ = 0.37. **C.** Group sizes: CT-VHC = 26, TR-VHC = 30, CT-CHX = 33, TR-CHX = 37. ANOVA of escape response during test session: F _(3, 122)_ = 6.07, *p* = 0.001, η^2^ = 0.13). Planned contrasts: CT-VHC vs TR-VHC, *t* _(122)_ = 3.01, *p* = 0.003, η^2^ = 0.14; CT-VHC vs CT-CHX, *t* _(122)_ = 1.86, *p* = 0.07, η^2^ = 0.06; CT-CHX vs TR-CHX, *t* _(122)_ = 2.32, *p* = 0.02, η^2^ = 0.07. Bayesian analysis: CT-VHC vs TR-VHC, BF_10_ = 5.48; CT-VHC vs CT-CHX, BF_10_ = 0.95; CT-CHX vs TR-CHX, BF_10_ = 3.59. \**p* < 0.05. \*\**p* < 0.01.

## Acknowledgements

This work was supported by National Agency of Scientific and Technological Promotion (PICT2013-0412 and PICT2016-0243) to MEP, and IBRO Return Home Fellowship to EM (GR037). Authors would like to thank Dr Gabriela Hermitte, Dr Jimena Berni and Dr Hans Crombag for their helpful comments on the manuscript.

